# BiofilmQ-HT: An integrated browser-based workflow for standardized analysis of high-throughput crystal violet-based biofilm assays

**DOI:** 10.64898/2026.09.04.749107

**Authors:** Atif Khan, Sruti Chandramouli, Hiren M. Joshi

## Abstract

The crystal violet (CV) microtiter plate assay is widely used to compare microbial biofilm formation because it is inexpensive, simple, and readily adapted to 96-well formats. Its high-throughput design enables multiple strains, growth conditions, and treatments to be tested in parallel, but increasing experimental complexity can make downstream data handling difficult. Plate layouts, absorbance measurements, blank correction, replicate analysis, and visualization are often managed using separate spreadsheets or software, requiring repeated transfer and organization of experimental information. This can compromise consistency in linking measurements with their corresponding conditions and controls. We developed BiofilmQ-HT, a standalone, browser-based workflow that integrates plate annotation with direct import of plate-reader data, condition-specific blank correction, replicate summarization, growth-normalized specific biofilm formation (SBF) analysis, percentage-inhibition analysis, visualization, and data export. The workflow was applied to three representative datasets: a comparison of biofilm formation in 25% and 100% media, an analysis of bacteriophage- and antibiotic-associated antibiofilm activity, and a screening of biofilm formation among natural microbial isolates. These applications demonstrate a common analytical framework for comparative biofilm analysis, antibiofilm screening, and high-throughput isolate screening while retaining experimental conditions and controls. By linking plate organization to quantitative analysis and reporting, BiofilmQ-HT provides a reproducible and traceable approach for processing condition-rich CV assay datasets and facilitates consistent comparisons across experimental groups. The utility therefore improves consistency in data handling without altering the underlying experimental assay or requiring changes to the established assay.

## 1. Introduction

Biofilms are structured microbial communities in which cells interact with surfaces and with one another within a self-produced extracellular matrix. This mode of growth is associated with spatial heterogeneity, altered physiology, and increased tolerance to environmental and antimicrobial stresses, making quantitative assessment of biofilm formation relevant across environmental, industrial, and biomedical microbiology [1, 2].

The crystal violet (CV) microtiter plate assay remains one of the most accessible approaches for comparative biofilm analysis. Its low cost, simple endpoint measurement, and compatibility with 96-well plates make it useful for testing multiple strains, growth conditions, and treatments in parallel. The assay, however, measures retained stain as a proxy for attached biomass, and the numerical output depends on experimental and analytical choices, including staining and solubilization procedures, measurement wavelength, background correction, and normalization. Recent work has emphasized the need for greater harmonization of CV-based measurements across laboratories [3–7]

A less visible source of variability occurs after the plate-reader measurement. In many laboratories, plate layouts are maintained separately from raw absorbance files, blank correction is performed through manually edited spreadsheets, replicate wells are summarized with laboratory-specific formulas, and plots or heatmaps are generated in separate programs. Semi-automated workflows have addressed parts of this problem, but practical analyses still require users to navigate between experimental metadata and analytical outputs [8]. When a plate contains several media, strains, treatment combinations, or controls, this separation makes it easier to apply the wrong blank, mislabel a condition, or lose the relationship between a well and its experimental context.

We developed BiofilmQ-HT to provide a simple analytical layer between the microplate reader and biological interpretation. The workflow retains the 96-well plate map during analysis and combines plate annotation, direct data import, condition-specific blank correction, replicate summarization, growth-normalized specific biofilm formation (SBF) analysis, percentage-inhibition analysis, visualization, and export. The purpose is not to replace the CV assay or impose a universal biological classification scheme, but to make routine data processing more explicit, reproducible, and less dependent on individually maintained spreadsheets. Three representative datasets are used to illustrate the workflow.

## 2. BiofilmQ-HT workflow

### 2.1. Design and implementation

BiofilmQ-HT is implemented as a client-side web application using HTML, CSS, and JavaScript and can be distributed as a standalone HTML file. The principal calculations are performed locally in the user’s browser rather than on a remote analysis server. This file-based design removes the need for a dedicated software installation or local server and allows a laboratory to retain and reuse a defined version of the workflow.

The analysis begins with an interactive 96-well plate map. Wells can be designated as Strain/Condition, Blank, Positive Ctrl, Negative Ctrl, or Empty, and each experimental well can be associated with a Media/Condition designation. Replicate wells are then grouped by the assigned strain or condition. In the datasets presented here, these were technical replicates and were used to describe within-plate measurement variability; they were not treated as independent biological replicates.

### 2.2. Data import and calculations

Reader-exported Excel or CSV files containing the expected 96-well measurement grid are imported directly into the workflow. For SBF analysis, OD600 and OD570 measurements are entered separately. OD600 is used as the growth-related measurement, and OD570 as the CV-associated measurement. For each experimental group, the workflow calculates the mean and standard deviation across replicate wells and associates the group with the blank that carries the corresponding condition designation.

For the current implementation, the blank-corrected SBF is calculated as:

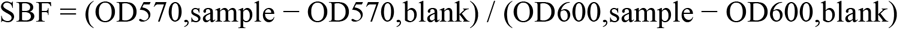

This growth-normalized calculation follows the SBF concept introduced by Niu and Gilbert, with blank correction applied to both OD570 and OD600 in the current implementation. Because published implementations differ in background correction and classification conventions, BiofilmQ-HT explicitly defines the formula used here to support reproducibility [9].

For antibiofilm experiments, percentage inhibition is calculated relative to the corresponding untreated Positive Ctrl after blank correction:

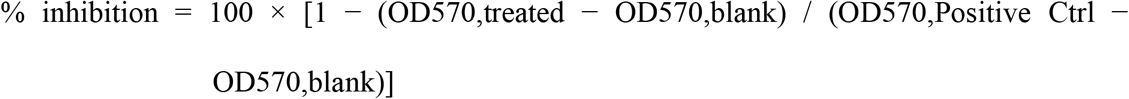

SBF and percentage-inhibition thresholds can be configured within the workflow. These categories are reporting aids rather than universal biological standards because CV responses depend on the organism, medium, staining procedure, measurement wavelength, and experimental design [3,6,7,10]. The configured threshold values are retained with the analysis and displayed in the corresponding output.

Automated processing does not eliminate the need for experimental quality control. Appropriate blanks and controls, complete well mapping, and biologically meaningful absorbance measurements should be verified before interpretation, particularly when blank correction produces values close to zero or when growth measurements are very low.

### 2.3. Visualization and export

Following the calculation, BiofilmQ-HT presents a summary dashboard, ranked plots, a detailed result table, and a 96-well heatmap. The result table retains group-level absorbance values, blank values, replicate variability, calculated SBF or percentage inhibition, and the assigned category. Results can be exported as CSV for downstream analysis and as PDF or image files for documentation, presentations, or manuscript preparation. Plate templates can also be saved and reloaded, allowing recurring experimental layouts to be reused without rebuilding the plate map.

**Figure 1.**
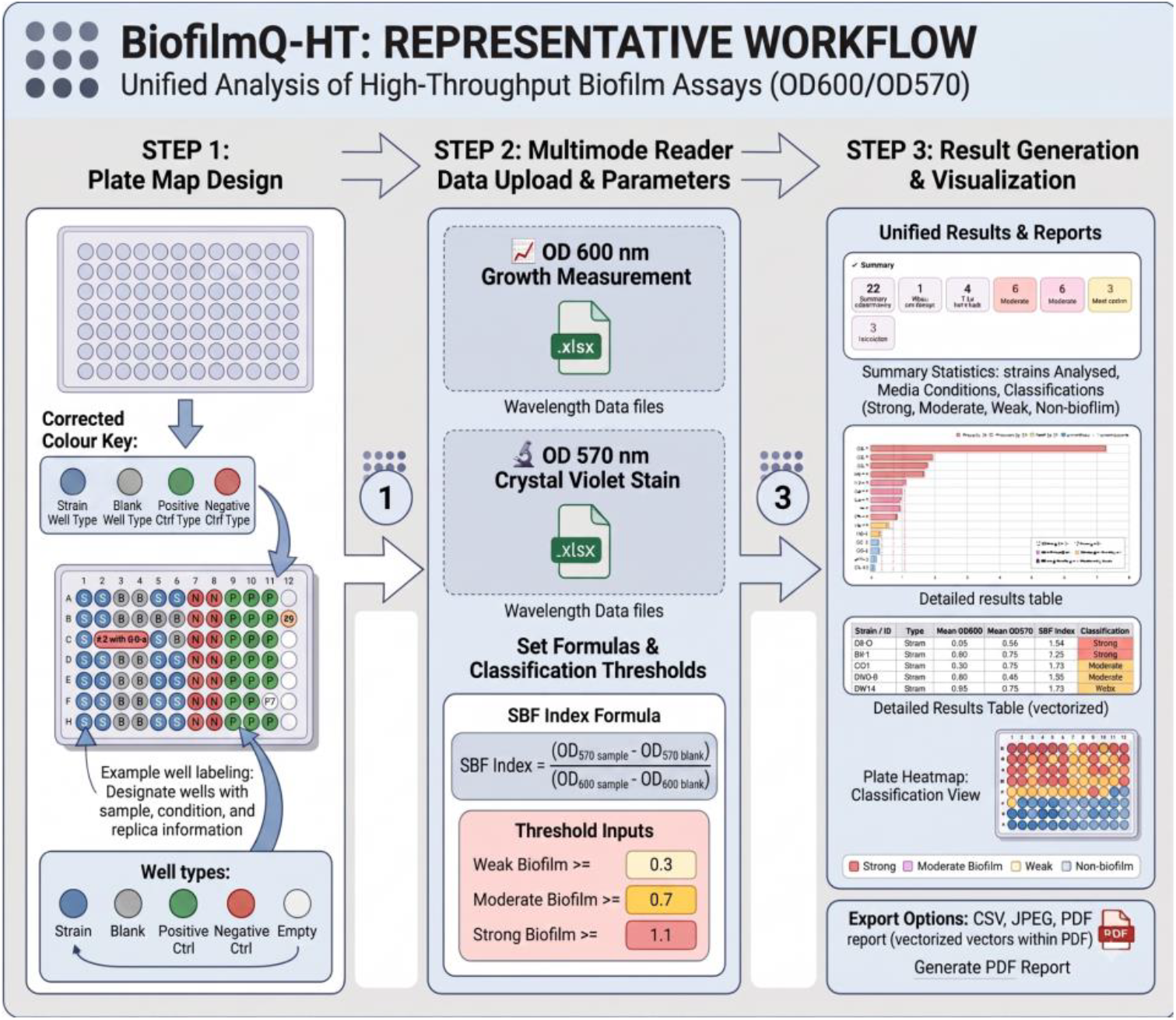
BiofilmQ-HT workflow. The workflow links 96-well plate annotation and condition metadata with OD600/OD570 data import, calculation, classification, visualization, and export.

Replicate wells are grouped by assigned strain or experimental condition. The workflow summarizes replicate measurements using the group means and standard deviation. In the datasets presented here, replicate wells represent technical replicates; they are therefore used to characterize within-plate measurement variability and are not treated as independent biological replicates.

## 3. Representative applications

### 3.1. Comparison of biofilm formation across growth-media conditions

BiofilmQ-HT was first applied to a dataset containing 15 microbial strains or conditions tested in parallel in 25% and 100% LB medium. Separate blank wells were included for each medium. The mean blank values were 0.1230 at OD570 and 0.0427 at OD600 for 25% LB, compared with 0.1489 at OD570 and 0.0509 at OD600 for 100% LB. SBF values varied among strains and between media. SS6 D7 and CT-6 had the highest SBF values in 25% LB (6.364 for both). In 100% LB, SS6 D7 remained a strong biofilm former (SBF 2.224), whereas CT-6 had an SBF of 1.243. FRP 6 had SBF values of 1.245 and 0.995 in 25% and 100% LB, respectively, while CT-7 showed values of 1.164 and 1.325 (the complete data set is provided in the supplementary information or can be downloaded from GitHub).

The principal point of this example is analytical rather than biological. Because the medium identity was retained in the plate map, each group was associated with the appropriate blank during calculation. The resulting output makes strain-specific differences between media visible without requiring manual reassignment of background values. The example illustrates how the workflow preserves experimental context.

**Figure 2.**
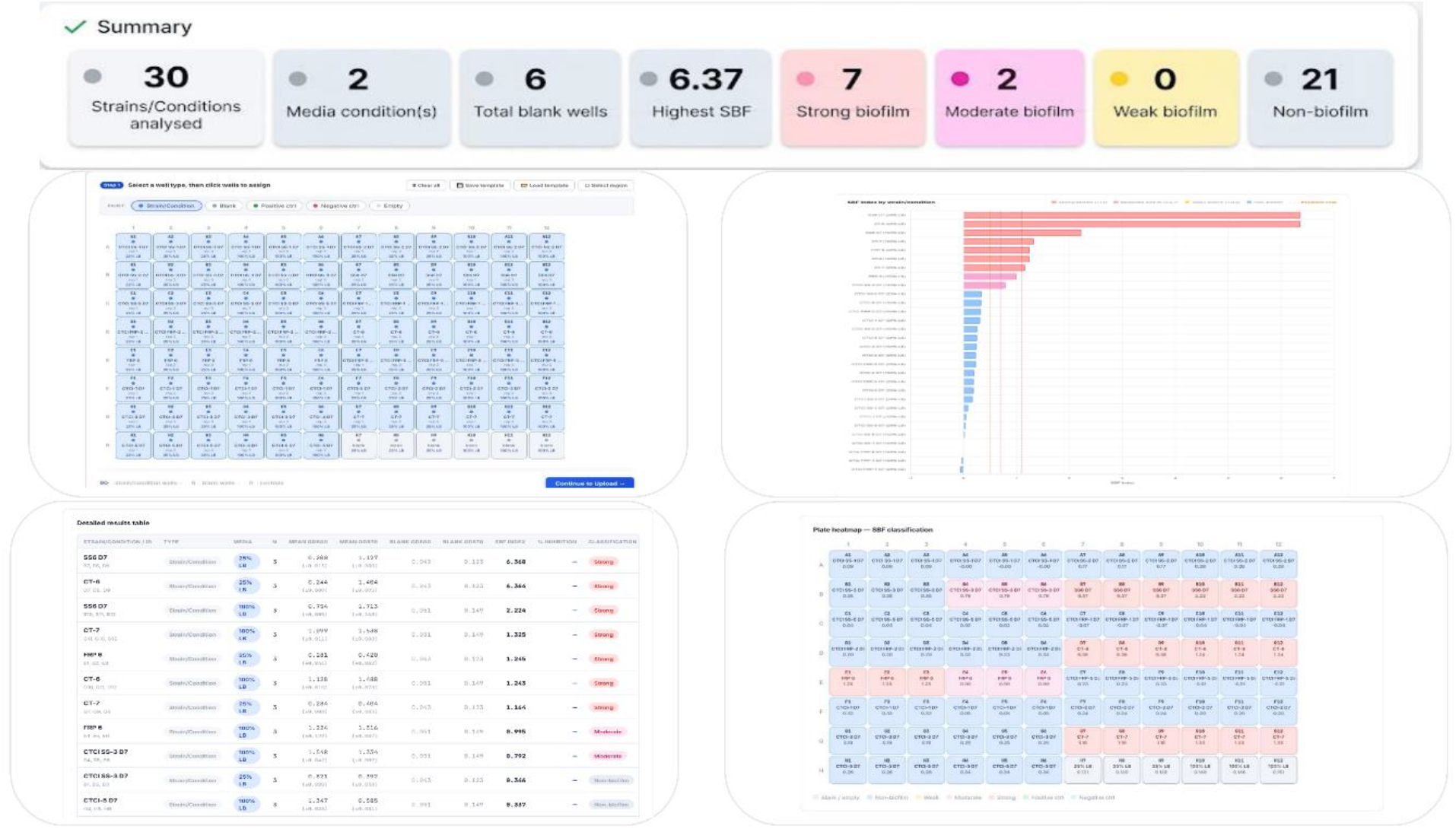
BiofilmQ-HT output for comparison of biofilm formation across growth-media conditions. The interface combines the plate map, ranked SBF values, group-level results, and plate-level classification.

### 3.2. Analysis of bacteriophage- and antibiotic-associated antibiofilm activity

The antibiofilm module was evaluated using an OD570 dataset containing an untreated Positive Ctrl, individual bacteriophage treatments, antibiotic treatments, and antibiotic-phage combinations. The untreated Positive Ctrl had a mean OD570 of 1.362 ± 0.034, and the blank had a mean OD570 of 0.115, giving a blank-corrected control value of 1.247. Individual phage treatments produced calculated inhibition values of 79.3% for Phage 1, 80.0% for Phage 6, 98.7% for Phage 8, and 69.2% for Phage 3. Amp-50 and Amp-300 alone produced 2.5% and 9.1% inhibition, respectively. Several antibiotic–phage combinations produced calculated inhibition values between 78.9% and 99.5%, with the highest value observed for Amp-50 + Phage 1 & 6 (99.5%).

These outputs demonstrate that multiple treatment groups can be processed against a common untreated reference while retaining their positions on the experimental plate. They should be interpreted as assay-level estimates of inhibition. Because the example uses technical replicate wells, the data do not establish biological reproducibility, interaction effects, or pharmacological synergy between antibiotic and phage treatments. Demonstrating synergy would require an appropriate experimental design and statistical model based on independent biological replicates.

**Figure 3.**
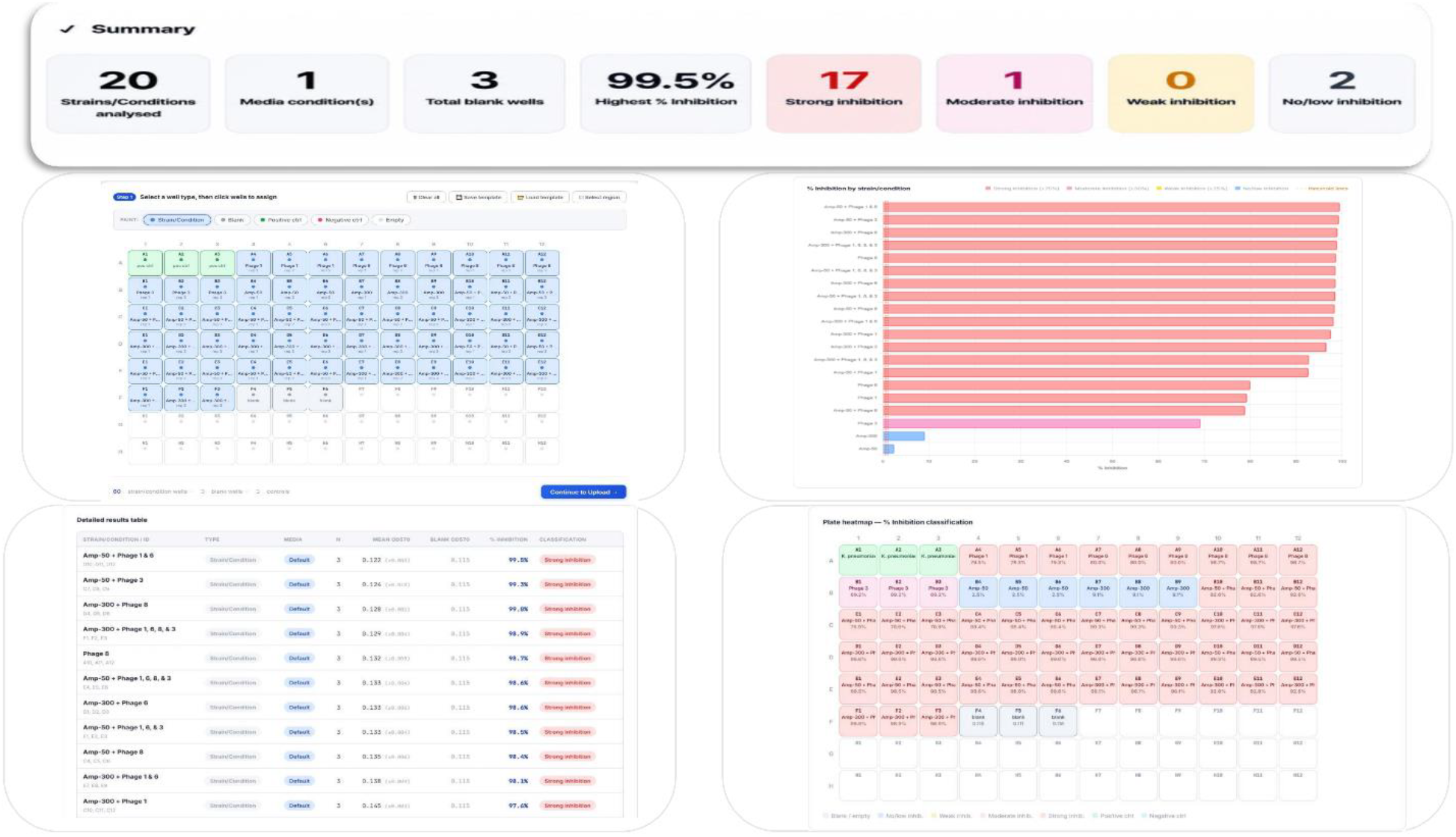
BiofilmQ-HT output for antibiofilm treatment analysis. Percentage inhibition is calculated from OD570 measurements relative to the untreated Positive Ctrl and displayed as ranked treatment results and a plate-level heatmap.

### 3.3. Screening microbial isolates for biofilm formation

The workflow was further applied to 23 microbial isolates, each represented by four technical replicate wells on the same plate. SBF values were calculated from group-level mean absorbances after blank correction. CB-5 had the highest SBF (7.343). Five additional isolates, SS 1 (1.919), CB-1 (1.794), FRP 1 (1.685), CB-15 (1.178), and CB-2 (1.175), were classified as strong biofilm formers using the configured threshold. CuNi 2 (0.964), CuNi 1 (0.930), CB-11 (0.924), Ti 1 (0.894), and CB-10 (0.848) were classified as moderate, while CB-16 was classified as weak. The remaining 11 isolates had SBF values below 0.5. Thus, 12 of 23 isolates were assigned to at least the weak-biofilm category under the configured classification scheme.

**Figure 4.**
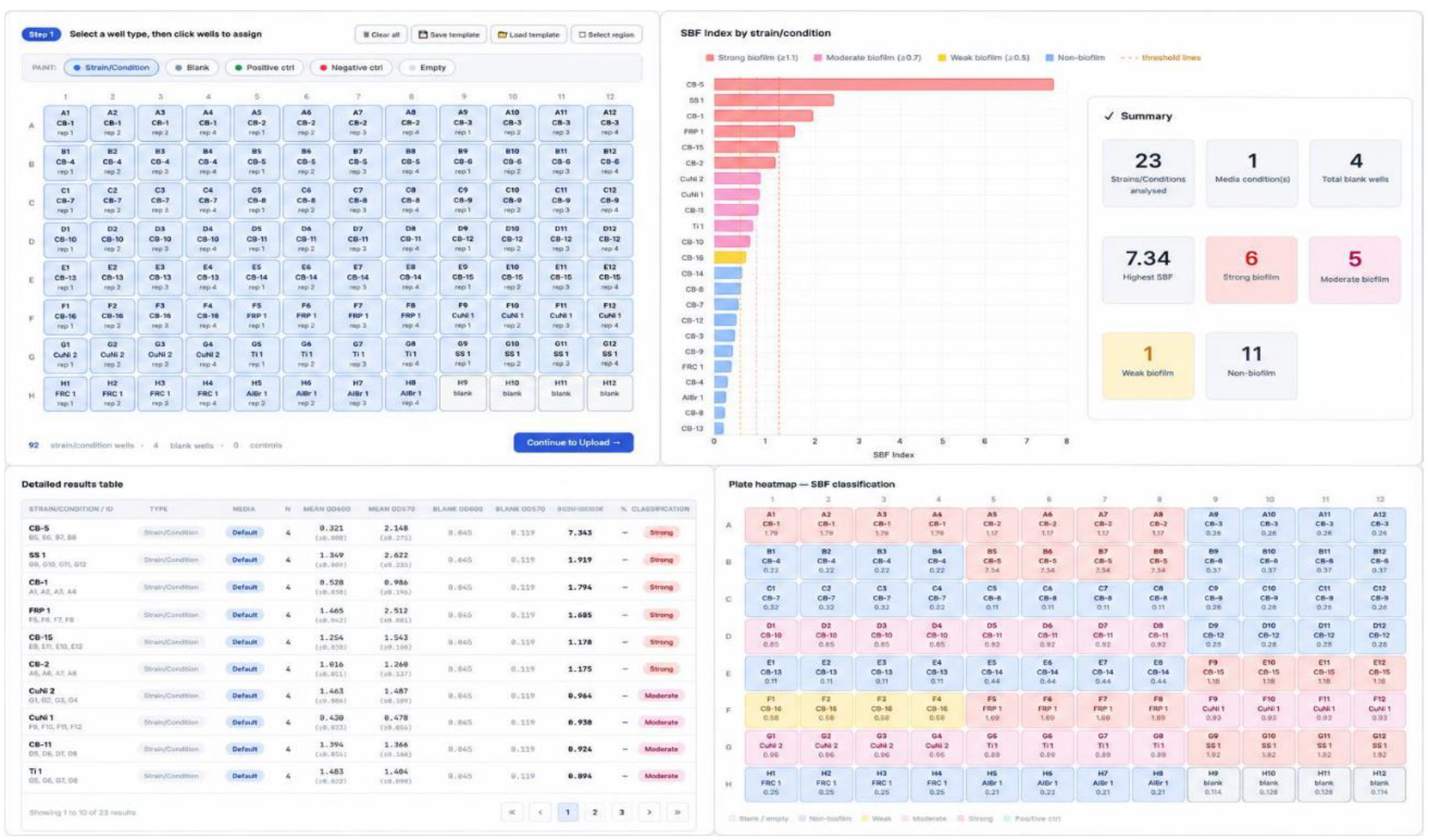
High-throughput screening of 23 microbial isolates using BiofilmQ-HT. Ranked SBF values are linked to the original plate layout and the configured classification thresholds.

The practical value of this application lies in the combination of quantitative ranking and plate-level classification. For screening studies, this allows the investigator to move from raw absorbance measurements to a structured list of candidates without manually rebuilding the plate layout in a second analytical environment. The categories remain operational labels and should be interpreted alongside the underlying SBF values and the experimental design.

## 4. Discussion

The CV assay is valuable precisely because it is accessible; a conventional plate reader and a 96-well plate can generate a large comparative dataset with limited instrumentation. The same simplicity, however, can encourage analytical workflows that depend on ad hoc spreadsheets and manually maintained plate maps. In biofilm research, this is more than a clerical issue. A blank, control, wavelength, or condition label is part of the experimental definition of a measurement. If that metadata becomes detached from the absorbance value, a technically correct calculation can still produce a biologically misleading result.

BiofilmQ-HT addresses this metadata problem by treating the plate map as part of the dataset rather than as a separate record. This is particularly useful when a plate contains multiple media or other background conditions. In the 25% versus 100% LB example, the workflow preserved the association between each group and its corresponding blank. The same principle applies to screening experiments in which treatment identity, concentration, or control type changes across the plate. The software therefore provides a reproducible path from experimental layout to calculated output without requiring the user to reconstruct that relationship during analysis.

The workflow also separates two concepts that are often conflated in routine CV studies: analytical standardization and biological standardization. BiofilmQ-HT can standardize how plate metadata are associated with measurements, how replicate groups are summarized, and how predefined calculations are applied. It cannot standardize the biology of the assay. Differences in organism, medium, incubation, staining, solubilization, measurement wavelength, and normalization strategy can alter the numerical response, and recent studies have highlighted the difficulty of comparing CV measurements across laboratories [3,6,7,10]. For this reason, the utility should be presented as a standardized data-processing framework rather than as a universal biofilm quantification standard.

The SBF implementation requires particular transparency. The SBF concept, introduced by Niu and Gilbert, normalizes CV-associated biomass to a growth metric, allowing attached biomass to be expressed relative to bacterial growth [9]. BiofilmQ-HT applies blank correction to both OD570 and OD600 in its current calculation. This operational definition is stated explicitly because published implementations differ in background correction and classification conventions. The software therefore reports the calculation along with the associated threshold settings, enabling the analytical procedure to be reproduced independently of the graphical interface.

The antibiofilm example illustrates another important boundary. The workflow can calculate inhibition values for phages, antibiotics, and combinations, but a high percentage of inhibition is not itself evidence of synergy. The same distinction applies to technical replicate variability: standard deviation among wells describes measurement dispersion within a plate, not biological reproducibility. BiofilmQ-HT is most useful when it makes these analytical distinctions visible rather than obscuring them behind automated categories.

A further strength is the browser-based, client-side architecture. Many laboratories do not need a large computational package for routine 96-well CV analysis, and a standalone HTML application can be easier to distribute, archive, and reuse than a software stack with multiple dependencies. The reusable plate template is also well suited to screening studies in which similar experimental layouts are repeated. These practical features are likely to matter most to biologists who need a transparent workflow rather than a general-purpose data-analysis platform.

The present version has clear limitations. It is focused on CV-based measurements and does not replace microscopy, viability assays, biomass calibration, or other orthogonal approaches. The three datasets demonstrate functionality but do not constitute a formal benchmark of processing time, error rate, or inter-user reproducibility. Future development could include explicit quality-control flags, broader plate-reader import formats, statistical analyses based on biological replicates, and additional biofilm readouts, while retaining the central link between experimental metadata and quantitative measurements [3–5].

## 5. Conclusions

BiofilmQ-HT provides a practical analytical layer for routine 96-well CV biofilm assays. By keeping the plate layout and experimental conditions linked to absorbance measurements, the workflow integrates blank correction, replicate summarization, SBF and percentage-inhibition calculations, classification, visualization, and export in a single browser-based environment. The representative datasets show that the same framework can accommodate media comparisons, antibiofilm treatment screens, and isolate collections. Its main contribution is not a new biofilm assay, but a more explicit and reproducible way to process the data generated by an established one. For laboratories performing repeated or multi-condition CV assays, this separation of biological measurement from ad hoc data handling can improve consistency without changing the underlying experimental workflow.

## Availability

BiofilmQ-HT is implemented in HTML, CSS, and JavaScript as a standalone browser-based application. The BiofilmQ-HT software, user manual, example datasets, and associated analysis files are available at https://github.com/phageforlifeai/BiofilmQ-HT (version 1.0.0).

